# The rugged adaptive landscape of an emerging plant RNA virus

**DOI:** 10.1101/006205

**Authors:** Jasna Lalić, Santiago F. Elena

## Abstract

RNA viruses are the main source of emerging infectious diseases owed to the evolutionary potential bestowed by their fast replication, large population sizes and high mutation and recombination rates. However, an equally important parameter, which is usually neglected, is the topography of the fitness landscape, that is, how many fitness maxima exist and how well connected they are, which determines the number of accessible evolutionary pathways. To address this question, we have reconstructed the fitness landscape describing the adaptation of *Tobacco etch potyvirus* to its new host, *Arabidopsis thaliana*. Fitness was measured for most of the genotypes in the landscape, showing the existence of peaks and holes. We found prevailing epistatic effects between mutations, with cases of reciprocal sign epistasis being common at latter stages. Therefore, results suggest that the landscape was rugged and holey, with several local fitness peaks and a very limited number of potential neutral paths. The viral genotype fixed at the end of the evolutionary process was not on the global fitness optima but stuck into a suboptimal peak.

## Introduction

Evolutionary dynamics of adaptation is a complex process depending on the complexity of the organism, as well as on many genetic, developmental, ecological and environmental factors. Sewall Wright’s metaphor of the fitness landscape (Wright 1932) pictures the process of adaptation as a surface in a multidimensional space that represents the relationship between mean fitness and the frequency of alleles within a population. Epistasis determines the topography of adaptive landscapes (Poelwijk et al. 2011; Whitlock et al. 1995; Wright 1932;) as well as the accessibility of adaptive pathways throughout the landscape; thus, it is central to understanding the course of evolution (Franke et al. 2011; Weinreich 2005; Welch and Waxman 2005). In absence of epistasis or in the case of magnitude epistasis, mutations give rise to either zero, positive or a negative fitness effect, regardless of the genetic background. This results in adaptive landscapes that are smooth and single peaked. In a smooth fitness landscape the evolution will always move uphill towards the single global optimum. In the case of sign epistasis, the sign of the fitness effect of a mutation depends on the genetic background, such that only a fraction of the total paths to the optimum are selectively accessible, i.e. contain only steps that confer a performance increase. Reciprocal sign epistasis, in which two mutations are individually deleterious but jointly advantageous, gives rise to rugged landscape with multiple local optima (i.e. peaks) (Poelwijk et al. 2011). The ruggedness of adaptive landscapes is critical to predict whether the evolving populations may reach the global optima or, by contrast, trough alternative evolutionary pathways, may get stuck into suboptimal fitness peaks (Kvitek and Sherlock 2011; Poelwijk et al. 2007, 2011; Whitlock et al. 1995; Weinreich 2005).

To reconstruct an empirical fitness landscape means to generate all possible genotypic intermediates that led to adaptation to a new environment and measure their fitness. Genotypes should bear all possible combinations of mutations fixed by the adapted genotype. Till now, simple empirical fitness landscapes have been characterized for either a single gene or promoter in bacteria (Chou et al. 2011; Dawid et al. 2010; Lunzer et al. 2005; Poelwijk et al. 2007; Weinreich et al. 2006), protozoa (Lozovsky et al. 2009), fungi (De Visser et al. 2009), and *Human immunodeficiency virus* type-1 (HIV-1) (Da Silva et al. 2010; Kouyos et al. 2012) being this the only virus for which such landscape has been inferred. The majority of these studies are based on two genotypes: the ancestral and the one well adapted to a given environment, differing mutually in a small set of known mutations. The largest empirical fitness landscape derived so far was for an evolved genotype that bared five mutations, that is, a total of 32 genotypes. In all cases, the landscape was rugged and the number of accessible evolutionary pathways driving to the highest fitness peak was limited.

Here, we aimed to construct the first empirical adaptive landscape for an emerging plant RNA virus. Our work is based on previous experimental evolution of host switching. Agudelo-Romero et al. (2008) simulated the emergence of *Tobacco etch virus* (TEV; genus *Potyvirus*, family Potyviridae) in a population of a partially susceptible host, *Arabidopsis thaliana* ecotype L*er*-0. *A. thaliana* belongs to family *Brassicaceae*, whereas TEV primary host species all come from a completely distinct family, the *Solanaceae*. Evolution of TEV on *A. thaliana* was done by serial passaging of the virus in several independent evolution lineages. After 17 serial passages, one of the evolved lineages, hereafter named TEV-*At*17, showed ca. 10-fold higher infectivity, 2-logs greater viral load, increased virulence and more severe symptoms in the new host compared to the ancestral virus (Agudelo-Romero et al. 2008). Transcriptomic analyses have shown that the evolved virus was able of evading host’s defense mechanisms by shutting down the expression of several defense pathways compared to its ancestor. During adaptation TEV-*At*17 fixed six mutations among which half were synonymous (table 1). Five different cistrons were affected by mutations (table 1). Agudelo-Romero et al. (2008) showed that symptoms development and increase in virus fitness were triggered by nonsynonymous mutation VPg/L2013F and that nonsynonymous mutations P3/A1047 V and 6K1/T1210M additively exacerbated the severity of symptoms in presence of mutation VPg/L2013F but had no effect in its absence. The VPg protein plays a central role in TEV production and spread (Jiang and Laliberté 2011). VPg is covalently attached to the 5’ end of the genome providing the hydroxyl residue that primes the synthesis of the complementary strand during transcription (Murphy et al. 1991). In addition, it triggers cap-induced translation via interaction with the eukaryotic translation initiation factor 4E (Robaglia and Caranta 2006) and it is involved in long-distance movement (Schaad et al. 1997).

**Table 1.**
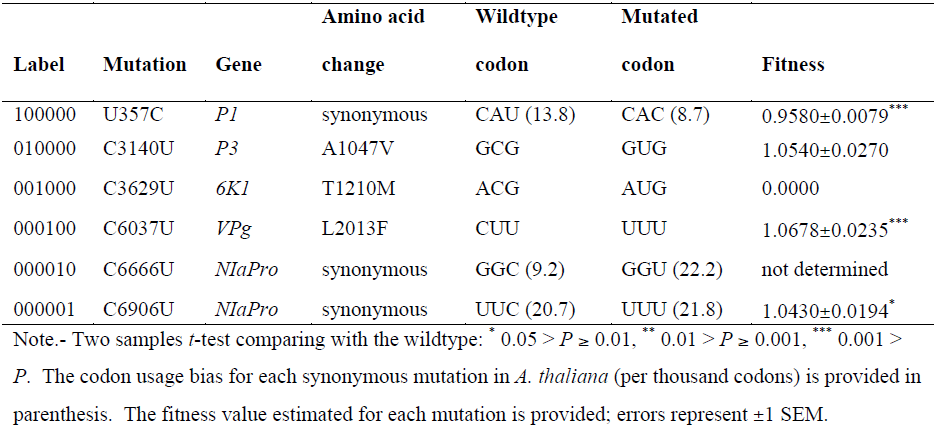
Mutations fixed in the *A. thaliana*-adapted virus TEV-*At*17.

| Label | Mutation | Gene | Amino acid | Wildtype | Mutated | Fitness |
| --- | --- | --- | --- | --- | --- | --- |
|  |  |  | change | codon | codon |  |
| 100000 | U357C | *P1* | synonymous | CAU (13.8) | CAC (8.7) | 0.9580±0.0079<sup>***</sup> |
| 010000 | C3140U | *P3* | A1047V | GCG | GUG | 1.0540±0.0270 |
| 001000 | C3629U | *6K1* | T1210M | ACG | AUG | 0.0000 |
| 000100 | C6037U | *VPg* | L2013F | CUU | UUU | 1.0678±0.0235<sup>***</sup> |
| 000010 | C6666U | *NlaPro* | synonymous | GGC (9.2) | GGU (22.2) | not determined |
| 000001 | C6906U | *NlaPro* | synonymous | UUC (20.7) | UUU (21.8) | 1.0430±0.0194<sup>*</sup> |

Note.- Two samples *t*-test comparing with the wildtype: <sup>\*</sup> 0.05 > *P* ≥ 0.01, <sup>\*\*</sup> 0.01 > *P* ≥ 0.001, <sup>\*\*\*</sup> 0.001 > *P*. The codon usage bias for each synonymous mutation in *A. thaliana* (per thousand codons) is provided in parenthesis. The fitness value estimated for each mutation is provided; errors represent ±1 SEM.

One way of representing fitness landscapes is in the form of a graph where each node corresponds to a specific genotype. Instead of representing genotypes in terms of nucleotides or amino acids, one can solely indicate absence or presence of a mutation at a given site, i.e. possible entries at each site are either 0 or 1, respectively, giving rise to a binary graph (fig. 1). With this notation, the wildtype genome is represented as 000000 and the TEV-*At*17 as 111111. Under the assumption of low mutation rates, the possible mutational steps are limited to single site substitutions, resulting in binary state space that has the topology of a Boolean hypercube. The arrows in fig. 1 show mutational transitions, connecting genotypes that mutually differ in a single position.

**Fig. 1.**
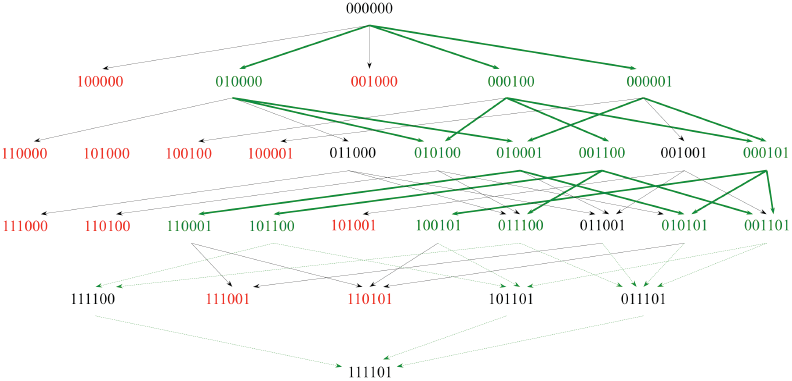
Empirical fitness landscape of size *n* = 5 for the evolution of TEV in *A. thaliana* L*er*-0. The presence/absence of a given mutation is indicated by 1/0, respectively. Genotypes are ordered top-down starting from the ancestral wildtype virus (000000) and finishing with the genotype carrying five of the six mutations fixed in TEV-*At*17 (111101). Each row represents those genotypes with equal number of mutations. Arrows represent all possible mutational transitions between genotypes. Genotypes marked in red are those with fitness lower than the ancestral wildtype virus (including lethals). Genotypes indicated in green are significantly fitter than the ancestral wildtype virus. Solid green lines represent likely mutational pathways (always connecting beneficial genotypes), dashed green lines pathways that involve a genotype not significantly better than the ancestral (neutral).

In order to picture the accessible evolutionary pathways for TEV-*At*17 in *A. thaliana* L*er*-0, we constructed most possible intermediate genotypes of adaptation and quantified fitness of each genotype in *A. thaliana* L*er*-0 relative to the 000000 wildtype genotype. One of the mutations (000010 in table 1) and all genotypes carrying this mutation were impossible to reproduce by site-directed mutagenesis because of unexpected problems with the stability of the corresponding mutant plasmids. Thus, the set of genotypes constructed consisted of 2^5^ = 32 combinations of five mutations fixed by TEV-*At*17 during its adaptation to the new host.

With this experiment we sought to answer important questions about an RNA virus adaptive landscape and its topography. First, we aimed to infer the prevalence of epistasis and characterize different types of epistasis that may be present in the architecture of fitness. Next, we question whether the differences in fitness are large relative to differences between genotypes, so that many changes are needed to obtain a high-fitness peak. In other words, how rugged is the landscape and how this topography influences the evolutionary potential of a virus population in the new host environment? If small genetic differences were associated to large differences in fitness, the landscape would be rugged. Conversely, if small differences in fitness were associated with large differences between genotypes, the landscape would be smooth.

## Results

### Synthetic lethal genotypes generate holes in the landscape

The very low efficiency of infection of *A. thaliana* L*er*-0 with in vitro transcribed viral RNAs generates the necessity of producing infectious viral particles in a susceptible host, usually the primary one *N. tabacum*, which are then used to inoculate arabidopsis. This is because the low efficiency to infect this plant using *in vitro* transcribed RNAs. Mutant genotypes 110000, 101000, 100100, 100001, 111000, 110100, 101001, 111001, and 110101 were not viable in *N.tabacum* (fig. 1), representing cases of synthetic lethality, i.e. mutations that were viable by themselves result in a non-viable genotype when combined. Interestingly, all these synthetic lethal genotypes share synonymous mutation *P1*/U357C. Since lack of viability in *N. tabacum* precludes generating the viral particles needed to inoculate *A. thaliana* L*er*-0, we could not directly test whether these genotypes were also lethal in the new host or, by contrast, they may be viable and thus lethality in tobacco may generate artificial holes in the fitness landscape. In first instance, we tested whether the lethality might result from the incorporation of undesired mutations during the *in vitro* transcription. Thus, we repeated the *in vitro* transcription twice, independently, for each genotype. Results where the same in both experiments: genotypes were lethal in *N. tabacum*.

In second instance, we tested whether genotypes lethal in tobacco might not really be so, but instead, may have lost their ability to systemically infect tobacco plants. Herewith, it is important to notice that the typical necrotic spots and vein etching produced by the ancestral TEV were not caused by the evolved TEV-*At*17. We performed a diagnostic one-step RT-PCR for TEV in the inoculated leaf using primers that specifically amplify 334 nucleotides of a conserved region from the virus *NIb* gene (Lalić et al. 2010). The tests for local infection were all negative: none of the tobacco plants was infected with either of the putative lethal mutant viruses. Nonetheless, to rule out that some low level replication was still taking place in the tobacco inoculated leaves, we prepared sap from each inoculated tobacco leaf and used it to inoculate 12 *A. thaliana* L*er*-0 plants per virus genotype. The result was, again, a lack of infection of the new host.

Thirdly, since we have previously shown the occurrence of antagonistic pleiotropy of mutational effects across alternative susceptible hosts (Lalić et al. 2011), we decided to test the viability of these viral genotypes on two other hosts highly susceptible to TEV infection: *Nicotiana benthamiana* and *Datura stramonium*. Inoculation of batches of each plant species with 5 µg of RNA produced by *in vitro* transcription of lethal genotypes within two independent blocks resulted in no infected plants. Therefore, we conclude that these nine genotypes must be truly lethal, thus generating holes in the adaptive landscape.

### Evolving TEV populations do not always move uphill

Overall, differences in relative fitness among genotypes exist in the new host (tables 1 and 2 and fig. 2; *χ*^*2*^ = 706.905, 31 d.f., *P* < 0.001), with 60.16% of the total observed variance among genotypes being explained by genetic factors. Eleven genotypes show significant differences in fitness compared with the ancestral TEV (two samples *t*-tests, FDR correction for multiple comparisons; *P* < 0.025). For these significant cases, relative fitness was widely variable. Indeed, two genotypes were worse than the ancestral TEV: genotype 001000 was lethal (notice that it was viable in tobacco) and genotype 100000 had a 4.20% fitness disadvantage. The other nine significant cases were all fitter than the ancestral TEV, with the smallest difference being observed for genotype 101100 (6.10%) and the largest one for genotype 010001 (12.01%).

**Fig. 2.**
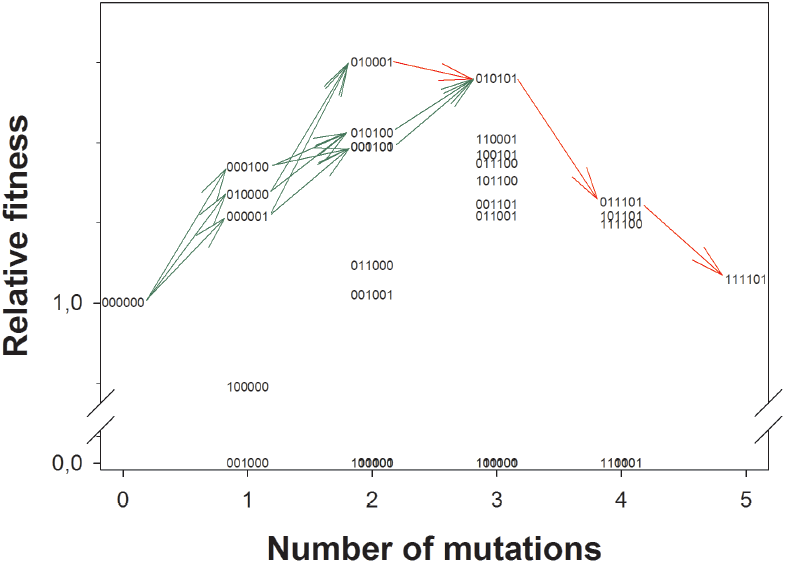
Relative fitness of each genotype as a function of the number of mutations that it carries. Genotypes are labeled according to the nomenclature used throughout the text. Genotypes 111100 and 111101 are cases of reciprocal sign epistasis. Arrows represent the most likely evolutionary pathways discussed in the text. Green arrows indicate mutations of beneficial effect on the genetic background where they are introduced. Red arrows indicate mutations of deleterious effect on the background where they appear.

**Fig. 3.**
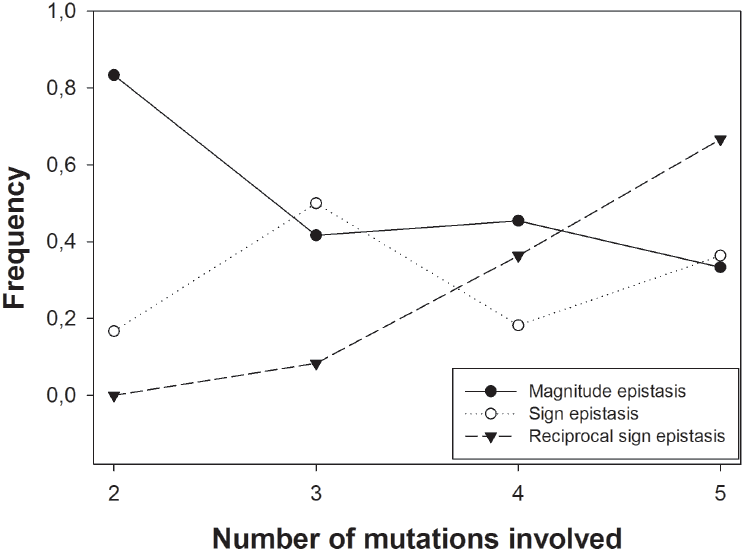
Effect of the number of mutations on the frequency of different types of epistatic interactions. Magnitude epistasis is more common for genotypes with low numbers of mutations while reciprocal sign epistasis becomes the most common type for large numbers of mutations.

Figure 2 illustrates the lack of monotonic increase in fitness from the ancestral TEV to the evolved TEV-*At*17 as the number of mutations introduced in the genome increased, supported by a non-significant Spearman’s correlation (*ρ* = −0.006, 30 d.f., *P* = 0.975). The maximum fitness value corresponds to genotype 010001 (carrying nonsynonymous mutation P3/A1047 V and synonymous mutation *NIaPro*/C6906U), followed by genotype 010101 (11.18% increase in relative fitness) that incorporates mutation VPg/L2013F to the 010001 genetic background. Therefore, TEV adaptation to *A. thaliana* is not conveniently explained by the simplistic picture of a population moving uphill in a smooth landscape. In the next sections we explore details of the empirical landscape topology that may help us understanding why TEV ends up in a suboptimal fitness peak.

A Tukey *post hoc* test classifies single mutants into three groups according to their fitness (*P* ≤ 0.002): the lethal genotype 001000, the low fitness genotype 100000, and a group formed by the other three single mutants, with genotype 000100 having the highest fitness. Double mutants could be classified into three non-overlapping groups according to their fitness (*P* ≤ 0.014): one group formed by the four synthetic lethals mentioned above, one formed by genotypes 001001 and 011000, of intermediate fitness, and one constituted by the rest of genotypes, among which 010001 has the highest fitness value. Likewise, triple mutants could be classified into two large groups (*P* ≤ 0.001): at the one side the three synthetic lethals and at the other side all viable genotypes, with 010101 being the one with the highest fitness among them. Finally, quadruple mutants could be classified into two non-overlapping groups (*P* ≤ 0.001): the two synthetic lethals, at the one side, and the three viable genotypes at the other side, with genotype 011101 having the highest fitness among them. Using this information, we can draw a likely scenario for the adaptive pathways followed by TEV during its adaptation to *A. thaliana*. The first mutation fixed was 010000, 000100 or 000001. From either of these mutations, genotypes 010100, 010001 and 000101 were accessible, all having higher fitness (two-samples *t*-tests, 1-tailed *P* ≤ 0.026 in all cases) (fig. 2). For the next step, the only genotype that is accessible from all previous ones is 010101, representing fitness increases in two cases (though not significant: two-samples *t*-tests, 1-tailed *P* ≥ 0.142) but fitness decline from 010001 (again, not significant: two-samples *t*-test, 1-tailed *P* = 0.189) (fig. 2). From there, evolution proceed downhill, with the next genotypes 011101 having lower fitness (but not significant: two-samples *t*-test, 1-tailed *P* = 0.072) and the final one 111101 even lower (in this case, marginally significant: two-samples *t*-test, 1-tailed *P* = 0.054) (red arrows in fig. 2).

### Epistasis and the ruggedness of TEV adaptive landscape

In the previous section we have shown that TEV-*At*17 does not necessarily represent the fittest genotype among others present in this empirical landscape, despite being the one fixed at the end of the evolution experiment. Indeed, genotypes 010001 and 010101 have significantly larger fitness values. Therefore, an explanation needs to be provided for this unexpected result. As we already discussed in the Introduction, the accessibility of adaptive pathways throughout the landscape depends on its ruggedness: the more rugged, the more constrained evolution may be and the more chances populations may get stuck into suboptimal peaks. A way to evaluate the ruggedness of this empirical landscape is to quantify the magnitude and type of epistasis among the five mutations.

The magnitude of epistasis among a set of *n* mutations was calculated as 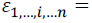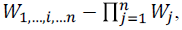 where *W*_*j*_ is the relative fitness of a genotype carrying mutation *j*, *W*_1,…,*i*,…*n*_ is the relative fitness of a genotype carrying mutations 1, …, *i*, …*n* and *ε*_1,…,*i*,…*n*_ is the epistasis coefficient between the 1, …, *i*, …*n* loci (Sanjuán and Elena 2006). The second term on the right-hand side of the equation corresponds to the expected fitness which, under the hypothesis of multiplicative independent effects, equals the observed fitness, resulting in *ε*_1,…,*i*,…*n*_ = 0. Deviations from the null hypothesis indicate antagonistic (positive) and synergistic (negative) epistasis, respectively. Epistasis coefficients calculated for each genotype carrying two or more mutations are listed in table 2. Observed epistasis values ranged from −1.125 (genotype 110001) to 1.078 (genotype 001100), with an average value of 0.417 ± 0.151 (±1 SD). This value was significantly positive (one-sample *t*-test: *t*_24_ = 2.760, *P* = 0.007), confirming previous observations that epistasis in TEV genome is predominantly positive (Lalić and Elena, 2012a). To evaluate the significance of epistasis in individual genotypes, we computed a *z* standard score for each of the 17 genotypes carrying at least two mutations. After applying the stringent FDR correction for multiple tests of the same null hypothesis, 16 genotypes showed a significant epistasis, five of them negative and 11 positive. All the 11 genotypes showing positive epistasis carried nonsynonymous mutation *6K1*/T1210M (001000) that, interestingly, was lethal by itself (table 1).

**Table 2.** Observed fitness and epistasis coefficients (*ε*) computed for each genotype carrying at least two mutations.

| Genotype | Fitness | Epistasis | *P* | Type of epistasis |
| --- | --- | --- | --- | --- |
| 110000 | 0 | -1.0097±0.0860 | < 0.0001 | Magnitude |
| 101000 | 0 | 0 |  |  |
| 100100 | 0 | -1.0230±0.0885 | < 0.0001 | Magnitude |
| 100001 | 0 | -0.9992±0.0670 | < 0.0001 | Magnitude |
| 011000 | 1.0186±0.0141 | 1.0186±0.0121 | < 0.0001 | Magnitude |
| 010100 | 1.0847±0.0187<sup>***</sup> | -0.0408±0.1309 | 0.3801 |  |
| 010001 | 1.1201±0.0164<sup>***</sup> | 0.0208±0.1128 | 0.3922 |  |
| 001100 | 1.0778±0.0215<sup>**</sup> | 1.0778±0.0148 | < 0.0001 | Sign |
| 001001 | 1.0038±0.0057 | 1.0038±0.0114 | < 0.0001 | Magnitude |
| 000101 | 1.0778±0.0220<sup>**</sup> | -0.0359±0.116 | 0.3802 |  |
| 111000 | 0 | 0 |  | Sign (100000 on 011000) |
| 110100 | 0 | -1.0782±0.1278 | < 0.0001 | Sign (100000 on 010100) |
| 110001 | 1.0816±0.0221<sup>**</sup> | 0.0284±0.1117 | 0.3862 | Sign (000001 on 110000, 010000 on 100001) |
| 101100 | 1.0610±0.0175<sup>**</sup> | 1.0610±0.0146 | < 0.0001 | Magnitude (000100 on 101000, 100000 on 001100), Reciprocal sign (001000 on 100100) |
| 101001 | 0 | 0 |  |  |
| 100101 | 1.0737±0.0283<sup>*</sup> | 0.0068±0.1145 | 0.3982 | Sign (000001 on 100100, 000100 on 100001) |
| 011100 | 1.0696±0.0304<sup>*</sup> | 1.0696±0.0147 | < 0.0001 | Magnitude (001000 on 010100) |
| 001101 | 1.0491±0.0288 | 1.0491±0.0150 | < 0.0001 | Magnitude (001000 on 000101) |
| 111100 | 1.0393±0.0212 | 1.0393±0.0129 | < 0.0001 | Magnitude (000100 on 111000, 100000 on 011100),<br>Reciprocal sign (001000 on 110100, 010000 on 101100) |
| 111001 | 0 | 0 |  | Sign (100000 on 011001) |
| 110101 | 0 | -1.1246±0.1504 | < 0.0001 | Sign (100000 on 010101), Reciprocal sign (000100 on 110001, 010000 on 010101) |
| 101101 | 1.0433±0.0300 | 1.0433±0.0153 | < 0.0001 | Magnitude (000100 on 101001, 001000 on 100101) |
| 011101 | 1.0501±0.0272 | 1.0501±0.0140 | < 0.0001 | Magnitude (001000 on 010101) |
| 111101 | 1.0117±0.0101 | 1.0117±0.0119 | < 0.0001 | Magnitude (000100 on 111001), Reciprocal sign (001000 on 110101, 010000 on 101101) |

Note.- Two samples *t*-test comparing with the wildtype: \* 0.05 > *P* ≥ 0.01, \*\* 0.01 > *P* ≥ 0.001, \*\*\* 0.001 > *P*. Fitness errors represent ±1 SEM. Epistasis errors represent ±1 SD. *P*-values correspond to the *z*-score test of significance of each epistasis coefficient.

One may ask whether the magnitude of epistasis tends to reduce as the viral population adapts to its new host, as should be expected for a smooth, single-peaked landscape. The curvature of such landscape should become more convex as the population approaches the optimum (Martin et al. 2007). Contrary to this expectation, we found no significant association between the magnitude of epistasis and the number of mutations fixed, that is, the distance to the genotype fixed (Spearman’s *ρ* = 0.234, 24 d.f., *P* = 0.250). This constancy of the average epistasis value regardless the position in the fitness landscape provides evidence that multiple peaks should exist, generating a highly rugged adaptive landscape.

As already mentioned in the Introduction, a necessary condition for ruggedness in adaptive landscapes is the existence of reciprocal sign epistasis (Poelwijk et al. 2011). In our previous study on epistasis among pairs of random deleterious mutations in TEV genome, we found a significant enrichment for this type of interaction (Lalić and Elena, 2012a). The question now is whether this conclusion also holds for the reduced set of potentially beneficial mutations being studied here. Following the mathematical conditions given by Poelwijk et al. (2011), we evaluated whether those cases for which we estimated a significant epistasis coefficient also corresponded to sign or reciprocal sign epistasis. Poelwijk et al. (2011) developed expressions for pairs of mutations and did not contemplated higher-order epistasis. Therefore, to be able of applying their formalism to genotypes carrying more than two mutations, we decomposed each genotype in all possible pairs. For example, genotype 101100 could be constructed in three ways: inserting mutation 000100 into genetic background 101000, inserting mutation 001000 into genetic background 100100, and by inserting mutation 100000 into 001100, meaning that we can test for three cases of epistasis for this genotype. This pairwise decomposition of interactions generates 64 possible genetic combinations for which epistasis shall be tested. Twenty-nine of these cases result in significant epistasis. Thirteen out of the 29 possible cases correspond to magnitude epistasis (table 2). In broad sense, sign epistasis was the most common type (16 of the 29; table 2). For example, we found that nonsynonymous mutations 001000 (6K1/T1210M) and 000100 (VPg/L2013F) fulfilled the condition for sign epistasis in the genetic background 001100. Furthermore, seven of the cases of sign epistasis also fulfilled the mathematical condition for reciprocal sign epistasis (table 2). This abundance of cases of sign, and especially of reciprocal sign epistasis, is consistent with the notion of a highly rugged adaptive landscape. In the case of genotype 111101, the condition was fulfilled either when mutation 010000 (P3/A1047 V) was incorporated into the genotype 101101 or when mutation 001000 appears in the 110101 background. Mutation 010000 shows reciprocal sign epistasis when it appears in genetic backgrounds 101101, 100101 and 101100. Mutation 0010000 also shows reciprocal sign epistasis when it appears in genetic backgrounds 100100, 110100, and 110101. All these genotypes share mutations *P1*/U357C and VPg/L2013F. The final case of reciprocal sign epistasis is mutation *NIaPro*/C6666U in genetic background 110001.

Figure 3 illustrates how the frequency of each type of epistatic interactions varies with the number of mutations present in the genotype. Cases of magnitude epistasis are more frequent for among double mutants but decrease as the number of mutations increase. By contrast, cases of reciprocal sign epistasis significantly increase with the number of mutations in the genotypes (Spearman’s *ρ* = 1.000, 4 d.f., *P* < 0.001). Cases of sign epistasis are roughly constant and independent of the number of mutations.

Therefore, we conclude that the adaptive landscape was smoother at earlier steps of the adaptation walk but increased in ruggedness as the number of mutations increases.

### Discussion

Here we address empirically the fitness landscape of a positive sense plant RNA virus TEV-*At*17 adapted to a novel host *A. thaliana*. Our dataset consisted of 32 possible combinations of five (out of six) mutations fixed by the evolved TEV-*At*17. We measured the relative fitness of genotypes that were still viable in the primary ancestral host, tobacco, on the new host, arabidopsis. Our results support the idea that this small empirical fitness landscape is highly rugged and holey, with ruggedness increasing as consequence of pervasive sign and reciprocal sign epistasis arising as the number of mutations introduced in the genome grows up. Upon reaching global fitness optima, represented by genotypes 010001 or 010101, the viral population drifted into lower fitness peaks after fixing slightly deleterious mutations on these genetic backgrounds.

A major portion of our analyses was focused on characterizing epistasis, measured as the deviation from the expected null hypothesis of multiplicative mutational fitness effects. Epistasis was prevalent (61.54%) in our dataset and predominately positive (68.75%), meaning that the absolute effect of the second mutation is smaller than that of the first. Previously, we have found significant prevalence of positive epistatic effects among deleterious mutations in TEV measured in its primary host *N. tabacum* (Lalić and Elena 2012a) as well as across a panel of alternative susceptible hosts (Lalić and Elena 2012b). Thus, positive epistasis is common in TEV genome among both deleterious and beneficial mutations and for both primary and alternative hosts. Strong mutational effects give rise to positive epistasis (Desai et al. 2007; De Visser et al. 2003, 2011; Proulx and Phillips 2005;), thus indicating low genetic robustness of RNA virus genomes. Furthermore, our results show that the epistasis governs subsequent steps of adaptation, as reciprocal sign epistasis becomes more common as the number of fixed mutations increases (fig. 1).

Analogous studies that sought to address the contribution of epistasis among beneficial mutations in bacteria (Chou et al. 2011; Khan et al. 2011) and yeast (Kvitek and Sherlock 2011) found a predominance of negative epistasis that impeded the rate of adaptation. Negative epistasis has diminishing returns effect meaning that adaptive mutations of large individual effect result in a smaller fitness effect when they occur together. Diminishing returns fitness effects were firstly observed in the experiments of long-term evolution of *Escherichia coli* (Barrick et al. 2009; De Visser and Lenski 2002) within which the initial fitness improvement was fast but it rapidly decreased over time and remained low. However, out fitness data show no pattern of diminishing returns, but instead, the adaptation began with relatively small steps in fitness increase until the fixation of the third mutation, a part from which, the fitness (in quadruple and quintuple mutants) decreased (fig. 1).

Epistasis causes stochastic differences in the rank order of mutations, hence, directly influences the evolutionary trajectories of populations (Martínez et al. 2011). Recent works on the contribution of epistasis to the architecture of fitness of RNA viruses found significant ruggedness of fitness landscapes. Lalić and Elena (2012a) found prevalent contribution of epistasis, especially of reciprocal sign type, to the architecture of TEV fitness suggesting that the adaptive landscape of TEV in its primary host (tobacco) must be highly rugged. Simultaneously, Hinkley et al. (2011) confirmed the commonality and strength of epistasis in HIV-1 protease and reverse transcriptase. In a concomitant study, Kouyos et al. (2012) analyzed fitness landscapes derived from *in vitro* fitness measurements of HIV-1 and reported ruggedness of the HIV-1 adaptive landscape. Because different peaks can also differ in magnitude, the adaptive landscape could impose an additional constraint on the evolvability of the organism. Rugged fitness landscapes have highly negative consequence on the evolvability of a population. Since a rugged fitness landscape lacks large constant-fitness plateaus, a population can become confined to a region of genotype space in which it must wait for the occurrence of the advantageous mutations. Our data support these former observations. Figure 1 and fig. 2 reveal that fitness landscape of TEV was multi-peaked; pervasive sign and reciprocal sign epistases are in the basis of this ruggedness (Poelwijk et al. 2007). We conclude that the fitness landscape of TEV on *A. thaliana* contains multiple adaptive peaks. Several other empirical studies have reached the opposite conclusion of empirical adaptive landscapes being smooth and single-peaked. Weinreich (2005) explored fitness landscape lacking sign epistasis and found a significant effect between landscape membership in fitness rank-ordering and genetic constraints of genotypes given by their fitness value. Later on, Weinreich et al. (2006) explored the fitness landscape of the *E. coli* β-lactamase conferring resistance to cefotaxime due to five mutations and found that majority of the trajectories contained fitness decreasing or neutral steps. Betancourt (2010) found no evidence of sign epistasis between beneficial mutations sweeping in independent lineages of evolving RNA bacteriophages. Kvitek and Sherlock (2011) observed few deleterious mutations that hitchhiked along with one or more adaptive mutations in the evolved yeast clones.

Mutations are random with respect to their effect in improving the fitness. Evolution is driven either by natural selection of beneficial mutations or by stochastic fixation of selectively neutral or slightly deleterious mutations due to random genetic drift. Mutation at locus *P1*/U357C showed independent fitness effects in all *n*-tuple mutant states implying that it was not important target of natural selection, but instead, results as a by-product of genetic drift. In favor to this idea goes the codon usage of *A. thaliana* that was lower for the mutated codon in comparison to codon of the wildtype virus (table 1). The evidences we presented here about the commonplace of epistasis together with the genotypes’ fitness results suggest that the evolution of TEV within *A. thaliana* L*er*-0 have not occurred solely by natural selection of mutations that improved TEV fitness within a new host. A major role in this evolutionary event most probably played the genetic drift associated with serial passages, especially in the fixation of the last two non-beneficial mutations. In nature, virus populations experience bottlenecks during transmission and cell-to-cell movement. Thus, genetic drift is an ever-present source of stochastic variation in allele frequencies among virus populations so it cannot be disregarded. In fact, genetic drift can lead to decrease in the mean fitness of an asexual population due to the process known as Muller’s ratchet (Mueller 1964) which proceeds if all individuals with the minimum number of deleterious mutations are lost by chance. Thus, Muller’s ratchet can lead a population into extinction (Lynch et al. 1993). In the case of independent mutational fitness effects, the rate of fitness decline is constant, but if there is a positive epistasis between deleterious mutations, as the ratchet advances, the frequency of the best available genotype will increase, making its loss less probable (Kondrashov 1994). Consequently, synergistic epistasis can arrest the action of Muller’s ratchet and provide the survival of the population although with lower mean fitness (Kondrashov 1994).

It is important to highlight three weaknesses of our study. First, missing data on synonymous mutation *NIaPro*/C6666U (000010) precludes the full understanding of the processes driving the evolution of TEV-*At*17 as accessible pathways may exist that require the fixation of this mutation. Second, we cannot argue that all tested genotypes form part of the same continuous network, so that mutations arose in a successive order. It is possible that independent mutational events in a highly polymorphic viral population may contribute jointly (by complementation or by recombination) to fixation of the observed TEV-*At*17 consensus sequence. In such situation, two or more distinct networks may exist, with multiple starting points and converging into certain nodes. Finally, we measured fitness as a relative within-host accumulation and thus neglected other fitness components such as transmissibility and stability that may compensate for the replicative disadvantage of genotypes 011101 and 111101.

## Methods

### Virus genotypes

Virus genotypes were constructed by successive rounds of site-directed mutagenesis starting from template plasmid pMTEV (Bedoya and Daròs 2010) and using mutagenic primers with specific single-nucleotide mismatch and Phusion^®^ High-Fidelity DNA Polymerase (Finnzymes) following manufacturer’s manual. PCR mutagenesis profile consisted of 30 s denaturation at 98 °C, followed by 30 cycles of 10 s at 98 °C, 30 s at 60 °C and 3 min at 72 °C, ending with 10 min elongation at 72 °C. Next, the PCR-mutagenesis products were incubated with *Dpn*I (Fermentas) for 2 h at 37 °C in order to digest the methylated parental DNA template. *E. coli* DH5α electrocompetent cells were transformed with 2 µL of these reactions products and plated on LB agar supplied with 100 µg/mL ampicillin. Bacterial colonies were inoculated in 8 mL LB liquid medium supplied with 100 µg/mL ampicillin and grown for 16 h in an orbital shaker (37 °C, 225 rpm). Plasmid preparations were done using Pure Yield^TM^ Plasmid Maxiprep System (Promega) and following the manufacturer’s instructions. Incorporation of mutation was confirmed by sequencing a ca. 800 bp fragment circumventing the mutagenized nucleotide. The plasmid DNA was *Bgl*II linearized and i*n vitro* transcribed using mMESSAGE mMACHINE^®^ SP6 Kit (Ambion) as described in Carrasco et al. (2007) in order to obtain infectious RNA of each virus genotype.

### Plants inoculation experiments

Since *A. thaliana* is extremely hard to be infected with naked RNA, but easy with entire virions, we used *N. tabacum* for production of virus particles. Batches of eight-week old *N. tabacum* plants were inoculated with 5 µg of RNA of each viral genotype by abrasion of the third true leaf. Ten days post-inoculation (dpi), the whole infected plants were collected and pooled for each virus genotype. Next, plant tissue was frozen with liquid N_2_, homogenized using mortar and pestle and aliquoted in 1.5 mL tubes. Saps were prepared by adding 1 mL of 50 mM potassium phosphate buffer (pH 8.0) per gram of homogenized plant tissue. Next, the homogenate was centrifuged at 4 °C and 10000 g for 10 min and the upper liquid phase with 10% Carborundum served as sap inocula.

*A. thaliana* L*er*-0 plants were grown in a BSL-2 greenhouse at 25 °C and 16 h light period. Plants were inoculated at growth stage 3.5 regarding the scale of Boyes et al. (2001). Six plants per virus genotype per block were inoculated with extracts of virus genotypes. The inoculations were done in three independent blocks. Infection was determined by one step RT-PCR as described previously (Lalić et al. 2010). Infected whole plants were collected at 21 dpi. Total RNA was extracted and virus accumulation was quantified by RT-qPCR as described in Lalić et al. (2010).

### Fitness calculation

Virus accumulation *Q* (pg of TEV RNA per 100 ng of total plant RNA) quantified at *t* = 21 dpi were transformed into Malthusian growth rate per day according to 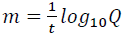 where (Lalić et al. 2011). Then, the relative fitness of each genotype (or just fitness, *W*) was calculated as 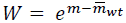, where 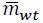 is the average Malthusian parameter estimated for the ancestral wildtype TEV.

Statistical tests were performed using SPSS version 20.

## Acknowledgements

We thank Francisca de la Iglesia for excellent technical support, José A. Daròs for helpful methodological advices and Jasper Franke for insightful comments. This work was supported by grant BFU2012-30805 from Spanish Dirección General de Investigación Científica y Técnica to S.F.E. J.L. was supported by a JAE-pre contract from CSIC.

